# ParGenes: a tool for massively parallel model selection and phylogenetic tree inference on thousands of genes

**DOI:** 10.1101/373449

**Authors:** Benoit Morel, Alexey M. Kozlov, Alexandros Stamatakis

## Abstract

**Motivation:** Coalescent- and reconciliation-based methods are now widely used to infer species phylogenies from genomic data. They typically use per-gene phylogenies as input, which requires conducting multiple individual tree inferences on a large set of multiple sequence alignments (MSAs). At present, no easy-to-use parallel tool for this task exists. Ad hoc scripts for this purpose do not only induce additional implementation overhead, but can also lead to poor resource utilization and long times-to-solution. We present ParGenes, a tool for simultaneously determining the best-fit model and inferring maximum likelihood (ML) phylogenies on thousands of independent MSAs using supercomputers.

**Results:** ParGenes executes common phylogenetic pipeline steps such as model-testing, ML inference(s), bootstrapping, and computation of branch support values via a *single* parallel program invocation. We evaluated ParGenes by inferring *>* 20, 000 phylogenetic gene trees with bootstrap support values from Ensembl Compara and VectorBase alignments in 28 hours on a cluster with 1024 nodes.

**Availability:** GNU GPL at https://github.com/BenoitMorel/ParGenes.

**Supplementary information:** Supplementary material is available at Bioinformatics online.

## 1 INTRODUCTION

The availability of genomic data for an increasing number of organisms allows to use thousands of genomic loci (henceforth: genes) to delineate evolutionary relationships between species. Species tree inference methods can be divided into supermatrix and supertree approaches. The former infer the species tree directly from a large concatenated MSA (*supermatrix*), whereas the latter infer individual per-gene trees which are then reconciled into a species phylogeny. Supermatrix methods are widely used due their simplicity and availability of efficient implementations (Nguyen *et al.*, 2015;Kozlov *et al.*, 2015). However, supertree inference methods gain popularity as they can model events such as incomplete lineage sorting (e.g.,Mirarab and Warnow (2015)), gene duplication and loss (e.g.,Arvestad *et al.* (2003)), as well as horizontal gene transfer (e.g.,Linz *et al.* (2007)).

As input, supertree methods typically require a set of per-gene trees (potentially also including bootstrap trees) that shall be reconciled (e.g., (Boussau *et al.*, 2012)) Inferring this set of per-gene trees using maximum likelihood (ML) methods is computationally intensive and requires the use of cluster computing resources.

While popular parallel tools for ML tree inference (e.g., RAxML (Stamatakis, 2014), IQ-TREE (Nguyen *et al.*, 2015)) can efficiently process large supermatrices, no dedicated parallel tool exists for inferring per-MSA trees on a large set of MSAs. In current studies users deploy ad hoc, and thus potentially error-prone, scripts for submitting each individual gene tree inference to a cluster as a single job. As cluster systems typically limit the number of sequential jobs a single user can execute in parallel, this can substantially increase the time-to-solution.

To this end, we have developed and made available a novel tool called ParGenes. It offers a simple command-line interface that allows to select the best-fit model, infer ML trees, and compute bootstrap support values on thousands of gene MSAs in a single MPI run. ParGenes relies on ModelTest-NG (https://github.com/ddarriba/modeltest) and RAxML-NG (Kozlov, 2018), to perform model selection and tree inference, respectively.

## 2 FEATURES

ParGenes encapsulates all per-gene calculations into one single MPI invocation. To improve load balance and decrease time-to-solution, ParGenes schedules per-gene inferences and allocates a *variable* number of cores to these inferences within its MPI runtime environment. In the following, we describe some of the key features.

### Simultaneous processing of MSAs

Unlike standard tools for ML inference, ParGenes analyzes multiple MSAs. Thus, the user needs to provide a directory containing all MSAs in PHYLIP or FASTA format. One can either specify global or MSA-specific options for both, RAxML-NG and ModelTest-NG. We pre-process each MSA, to check that the file is valid, compress it, save it in a binary file, and read its number of taxa and unique patterns.

### Model selection

ParGenes employs ModelTest-NG, a re-designed, substantially more efficient version of the widely used Modeltest tool (Posada and Crandall, 1998), to select the best-fit model of evolution for a given MSA. If model testing is enabled in ParGenes, it will first execute ModelTest-NG on each MSA, and then use the best-fit model for subsequent ML inferences.

### ML searches and bootstrapping

ParGenes schedules the per-MSA inference jobs that are executed using RAxML-NG (Kozlov, 2018). ParGenes allows to run multiple RAxML-NG tree searches per MSA from independent starting trees, which is recommended to better explore the tree search space. Then, it identifies the best-scoring ML tree for each gene. To increase job granularity and thereby improve load balance, each independent tree search is separately scheduled. ParGenes can also conduct a user-specified number of bootstrap (BS) inferences. It schedules independent tree inferences of BS replicates (10 BS replicates per job), and subsequently concatenates the resulting trees into one per-MSA BS tree file. Then, it runs RAxML-NG again to compute support values.

### Checkpointing

Since ParGenes runs are massively parallel and compute-intensive, it offers a checkpointing feature that allows for resuming ParGenes calculations (e.g., if program execution was interrupted due to typical cluster run-time limitations of 24 or 48 hrs).

### Estimating the optimal number of cores

Given the input MSAs, ParGenes can calculate an *a priori* estimate of the number of overall cores that will yield ‘good’ parallel efficiency. This is important, as it is difficult for users to set this value prior to running the analysis.

## 3 JOB SCHEDULING

ParGenes implements a scheduler that simultaneously executes independent jobs with a varying number of cores per job. A job is either a per-MSA RAxML-NG or ModelTest-NG run. We first outline the parallelization scheme, and then the scheduling strategy.

### Parallelization scheme

For the typical use case, the input data will contain thousands of independent (per-gene) MSAs with hundreds to a few thousand sites each. While standard tools like RAxML parallelize likelihood computations over MSA sites, ParGenes parallelizes the computations over the MSAs. Note that, the parallel efficiency of the RAxML parallelization is limited by MSA length (rule-of-thumb: 1,000 MSA sites per core). While most of input MSAs are small, their size exhibits substantial variance with respect to both, the number of taxa, *and* sites (Supp. Mat., Fig. 1). Therefore, inferring trees on large per-gene MSAs on a single core has two drawbacks. First, the MSA size might exceed the available main memory per core. Second, this can decrease parallel efficiency as a large job might take longer to complete than all other jobs (Supp. Mat., Fig. 2a). To this end, ParGenes allocates several cores for the largest jobs (MSAs) by invoking the respective multi-threaded RAxML-NG executable. For each MSA, ParGenes first calls RAxML-NG in parsing mode to obtain the recommended number of cores for optimal parallel efficiency via the fine-grained parallelization of the likelihood function in RAxML-NG (Stamatakis, 2015). The actual number of cores assigned to a job is then rounded down to the next power of two to simplify scheduling. We also assign twice the number of cores to the 5% MSAs with the largest number of taxa (Supp. Mat., Sect. 2).

### Scheduling strategy

ParGenes first sorts all jobs by (i) decreasing number of required cores and (ii) decreasing overall number of characters per MSA. As the number of cores per job (see Section 3) is always a power of two, ParGenes can always keep all cores busy, as long as there are jobs left to process. This works because the MSAs requiring the largest number of cores are scheduled first.

## 4 RESULTS

We evaluated ParGenes on two large empirical datasets obtained from Ensembl (Zerbino *et al.*, 2018) and VectorBase (Emrich *et al.*, 2015). They comprise 8, 880 and 12, 000 gene families, respectively. Executing the entire ParGenes pipeline on 1024 cores (model testing, ML tree search from 20 starting trees, bootstrapping analysis with 100 replicates) took 25 hours for the Ensembl dataset, and 3 hours for the VectorBase dataset. The VectorBase dataset required less time as its MSAs are smaller. In the supplement material, we show scalability results for different core counts.

## 5 CONCLUSIONS AND FUTURE WORK

We have presented an efficient parallel tool for comprehensive phylogenetic inference of gene trees on thousands of MSAs via a *single* MPI invocation. Apart from being flexible with respect to the inference options, ParGenes also yields ‘good’ parallel efficiency via appropriate scheduling mechanisms. We expect that ParGenes will contribute to increasing throughput times and productivity in gene-tree/species-tree reconciliation studies. Future directions entail the improvement of fault-tolerance mechanisms (e.g., core failures or single jobs failing for other reasons) and more accurate RAxML-NG runtime prediction approaches (e.g., machine learning).

## ACKNOWLEDGEMENT

This work was financially supported by the Klaus Tschira Foundation and by DFG grant STA 860/4-2. We are grateful to B. Bousseau and E. Tannier for providing the datasets.

